# State-dependent energy conversion produces degenerate dissipation in active actomyosin networks

**DOI:** 10.64898/2026.04.22.720181

**Authors:** Zachary Gao Sun, Juanjuan Zheng, A. Pasha Tabatabai, Joost J. Vlassak, Michael Murrell

**Affiliations:** Department of Physics, Yale University, 217 Prospect Street, New Haven, Connecticut 06511, USA; Systems Biology Institute, Yale University, 850 West Campus Drive, West Haven, Connecticut, 06516, USA; Integrated Graduate Program in Physical and Engineering Biology, Yale University, New Haven, Connecticut 06520, USA; John A. Paulson School of Engineering and Applied Sciences, Harvard University, Cambridge, MA, USA; Department of Biomedical Engineering, Yale University, 55 Prospect Street, New Haven, Connecticut 06511, USA

## Abstract

In non-equilibrium (active) systems, increased driving is commonly assumed to produce proportionally greater energy dissipation. Using picowatt-sensitive calorimetry, entropy-production analysis, rheology, and microscopy in reconstituted actomyosin networks, we show that this relationship breaks down as the material reorganizes under motor activity. Dissipation initially increases with myosin abundance but subsequently decreases despite continued network stiffening, indicating that energetic cost becomes regulated by the evolving mechanical state rather than actuator abundance alone. Comparing catch-bond (α-actinin) and slip-bond (fascin) crosslinked networks reveals that bond mechanics shift the critical motor concentration at which this transition occurs, marking the onset of state-dependent energy conversion. Although these networks generate distinct active stresses and mechanical states, they can exhibit comparable dissipation rates, revealing that state-dependent energy conversion can produce degenerate dissipation, whereby mechanically distinct states exhibit similar energetic costs. These findings demonstrate that dissipation in active materials is governed not only by the magnitude of driving but also by the mechanical state that emerges in response to that driving, providing an experimental example of state-dependent energy conversion far from equilibrium.

## Introduction

In non-equilibrium systems, increased driving is commonly assumed to amplify energy dissipation, reflecting intuition inherited from linear response theory and irreversible thermodynamics^1–6^. This intuition links activity, force generation, and energetic cost, in that stronger driving is expected to generate larger fluxes and greater heat loss. As a result, efficiency in systems ranging from molecular machines to active solids is often understood under the assumption that enhanced activity is accompanied by increased dissipation. Whether this assumption remains valid when systems are driven far from thermodynamic equilibrium, however, remains largely unexplored.

Linear response theory predicts that fluxes scale proportionally with their conjugate forces and that entropy production increases monotonically with driving^1–6^. In this regime, dissipation is determined by transport coefficients that are independent of the applied force, and added input is converted directly into heat through Onsager reciprocity^2^. This framework has been remarkably successful in describing transport^7^ and pattern formation in passive matter^8^, and is often implicitly extended to active systems^9–12^. Active materials, however, operate far from equilibrium, where driving does not merely induce motion but can alter the material itself^9,10,13^. In cytoskeletal networks, for example, internally generated stresses modify connectivity, stiffness, and force transmission, introducing feedback between mechanical organization and energy consumption^14–16^. Among these feedbacks are load-dependent molecular interactions^17–20^, which endow cytoskeletal materials with the ability to transition between solid-like and fluid-like states through motor-generated stresses transmitted by filamentous networks^17,21–24^. In such systems, dissipation may no longer be determined solely by the magnitude of driving, but also by how the material reorganizes and redistributes energy across its internal degrees of freedom. Indeed, recent experiments demonstrated that F-actin architecture can directly regulate the conversion of chemical energy into mechanical work, indicating that network organization can influence energy conversion itself ^14,25^. Directly testing this possibility has been challenging because energetic measurements are rarely combined with simultaneous characterization of material organization and mechanics. Previous work using stochastic thermodynamics and filament-scale fluctuations reported a non-monotonic dependence of entropy production rate on myosin abundance in quasi-two-dimensional actomyosin networks ^26^. Because active systems operate far from thermal equilibrium and generally lack a single effective temperature, entropy production inferred from stochastic fluctuations need not directly be equivalent to calorimetric heat dissipation. Thus, although previous studies showed that nonequilibrium activity can vary non-monotonically with motor abundance, it is unclear whether energetic dissipation is governed primarily by the magnitude of motor-driven chemical forcing or by the evolving architecture and mechanical state of the material. Distinguishing between these possibilities is essential for understanding how active materials convert chemical energy into mechanical work and heat.

In active materials, the quantity that is experimentally controlled is not necessarily identical to the quantity that ultimately transmits force or dissipates energy^27^. For example, increasing motor abundance may alter network architecture, motor loading, and ATPase kinetics, such that actuator abundance, active stress, and dissipation become distinct quantities. Therefore, throughout this work, we use myosin concentration as the experimentally imposed control parameter (“driving”), while exploring the potential that the resulting active stress emerges from the coupled motor-network dynamics and is not prescribed independently.

Here we show that increased driving does not necessarily amplify dissipation in active materials. Using the actomyosin cytoskeleton as a model non-equilibrium material^22–24,28–30^, we systematically vary myosin abundance and directly measure energy dissipation, mechanical response, and network dynamics. In parallel, we assemble the networks with catch-bond (α-actinin) and slip-bond (fascin) crosslinkers to alter the mechanical properties (state) of the system. Catch bonds strengthen under tensile load^21,31–34^, whereas slip bonds weaken with increasing tension^35,36^. These distinct force-dependent interactions provide a means to tune how active forces are transmitted through the network and define the mechanical state. By combining confocal fluorescence microscopy, entropy-production analysis^26^, picowatt-sensitive calorimetry^16^, and rheology, we find that dissipation exhibits a non-monotonic dependence on motor abundance. Networks occupying distinct mechanical states can exhibit similar dissipation rates, revealing a degeneracy between energetic cost and mechanical organization. Thus, motor abundance does not merely set the energetic cost; it reorganizes the network and thereby changes how ATP-derived energy is partitioned into work, structural remodeling, and heat. These results demonstrate that far-from-equilibrium active materials can decouple energetic cost from actuator abundance, and that energetic dissipation is not a unique indicator of mechanical state.

## Results

In this work, we assemble model actomyosin networks in two configurations ^37^. First, we assemble a quasi-2D network, amenable for fluorescence imaging and the quantification of filament dynamics ^22,37,38^. From this, we quantify how system state is changed by composition. Second, we assemble a 3D network, more amenable to rheology^39–41^ and calorimetry^42^ (Methods). With this assay, we identify mechanical and energetic signatures of non-equilibrium driving. To this end, integrating 3D imaging with pico-calorimetry allows direct comparison of network local deformation and heat dissipation, while rheological measurements provide complementary mechanical characterization. The molar ratios of proteins (e.g., crosslinker-to-actin and myosin-to-actin) are identical in the 2D and 3D assays, although the use of a crowding agent in the 2D system results in higher effective surface protein concentrations compared to bulk concentrations in 3D networks that are without crowding agent.

### Catch- and slip-bond crosslinkers produce distinct active mechanical states (*X*)

We reconstitute a quasi-two-dimensional active actomyosin network by confining pre-polymerized actin filaments to a supported lipid bilayer using 0.25% methylcellulose and crosslinking the network with either fascin or α-actinin. Fascin exhibits slip-bond behavior, whereas α-actinin forms catch bonds^22,37,38,43,44^ (Fig. 1a). Skeletal muscle myosin II is subsequently introduced to generate active contractile stresses within the network.

**Fig. 1.**
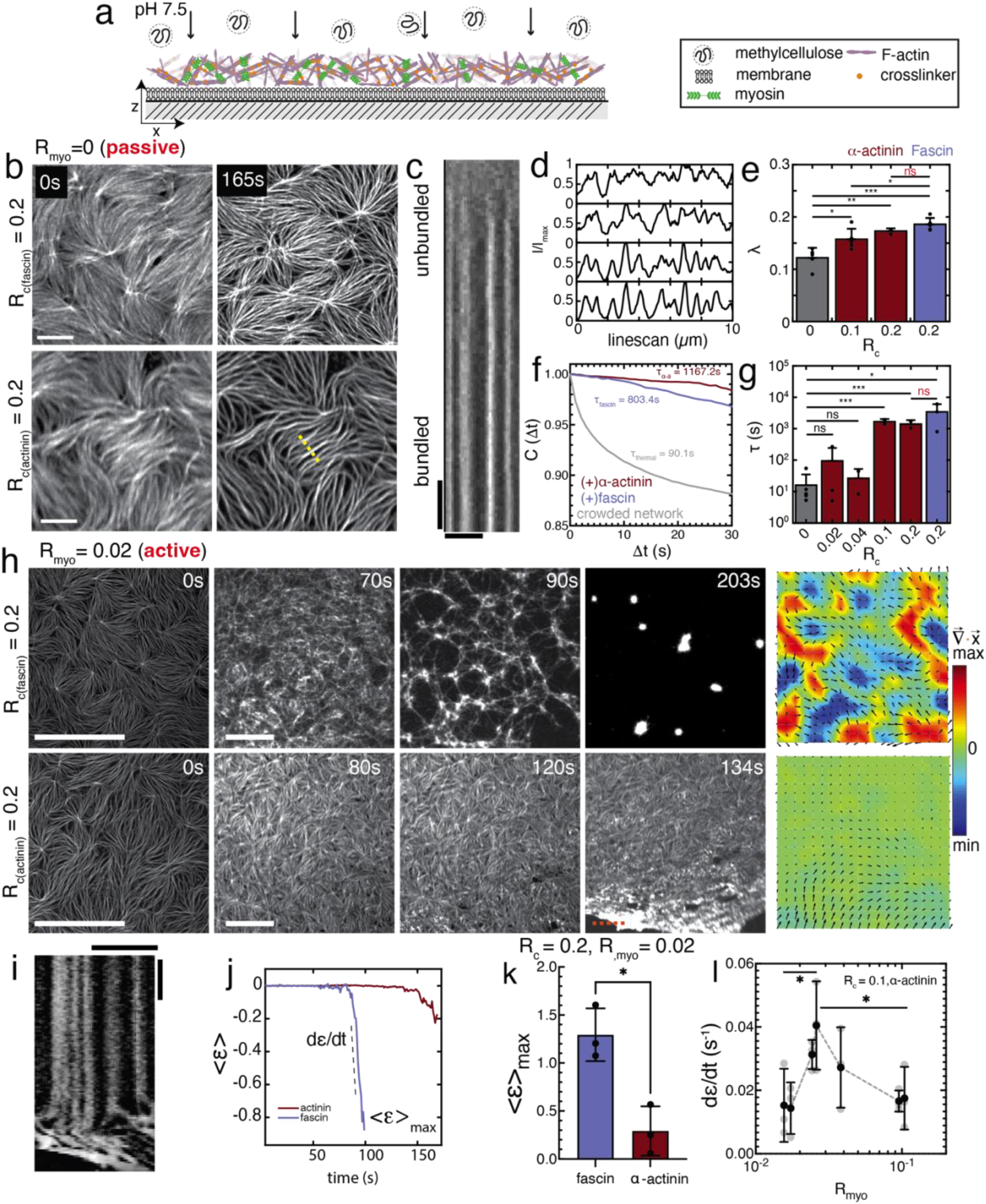
F-actin crosslinking determines distinct active material properties. (a) Schematic of pseudo-2D actomyosin network crowded by Methylcellulose on the lipid bilayer. (b) Confocal fluorescence microscopy image of pseudo-2D actomyosin network crosslinked by fascin (top) and α-actinin (bottom) on a supported lipid bilayer. Left is before crosslinking, and right is post crosslinker addition. Scalebar is 5 µm. (c) Kymograph of the yellow dashed line in b), scalebars are 5µm (horizontal) and 10s (vertical). (d) Exemplary linescan intensity (normalized) for conditions in e. (e) Bundling metric λ of fascin and α-actinin crosslinked networks as well as F-actin network without crosslinkers. N = 3 independent experiments for each condition. Error bars are the s.d. of the mean. (f) Exemplary autocorrelation function of actin network fluctuations as function of lag time Δ*t*. (g) Characteristic time *τ* for different conditions. N = 5,3,3,3,3,3 independent experiments respectively. Error bars are the s.d. of the mean. (h) Confocal fluorescence microscopy image of 2D actomyosin network crosslinked by fascin (top) and α-actinin (bottom) deformed and contracted under myosin active stress over time. Scalebars are 20 µm. Heatmaps show accumulative strain of the final frame. Quiver plot overlay shows the instantaneous velocity. (i) Kymograph of α-actinin network rupture (dashed red line in h). Scalebars are 10µm and 10s. (j) Mean strain < *ε* > of the network during deformation caused by myosin induced active stress over time. The slope of the strain curve is the strain rate 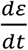. (k) Maximum mean strain < *ε* >_*max*_ of fascin and α-actinin crosslinked networks at R_c_ = 0.2. N = 3 independent experiments for each condition. Error bars are the s.d. of the mean. Welch’s t-test is performed. *p*_*fas*−*aa*_ = 0.0101. (l) Strain rate 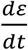 of the network crosslinked by α-actinin at R_c_ = 0.1 at various myosin concentrations. N = 3 independent experiments for all concentrations. Error bars are the s.d. of the mean. Standard linear model with least squares regression is performed. *p*_*up*_ = 0.0367, *p*_*down*_ = 0.0425.

The thermodynamic driving force for ATP hydrolysis, Δ*μ*_*ATP*_, is maintained approximately constant by the ATP regeneration system and therefore does not serve as the experimental control parameter. Instead, we vary the abundance of ATP-consuming myosin motors (*R*_*myo*_), thereby altering the number of mechanochemical actuators operating under an approximately fixed chemical potential difference.

At high crosslinker density R_c_ > 0.2, α-actinin- and fascin-crosslinked networks are morphologically similar when observed by optical microscopy (Fig. 1b–d). Both crosslinkers organize F-actin into bundled architectures, producing networks that are difficult to distinguish based on static morphology alone (Fig. 1d–e). Under thermal fluctuations, the two network types also exhibit comparable relaxation dynamics and characteristic relaxation times, suggesting similar passive mechanical properties (Fig. 1f–g).

Despite these similarities, the networks respond differently once active stresses are generated by myosin motors. Fascin-crosslinked networks contract into localized aster-like structures (Fig. 1h, Supplementary Movie 1) ^45^, whereas α-actinin-crosslinked networks undergo large-scale deformation and frequently fail through crack formation^39^ (Fig. 1h–i, Supplementary Movie 1). These distinct contraction modes are accompanied by significant differences in both the magnitude of accumulated strain (*ε*) and the time required for material failure (Fig. 1j–k).

To quantify differences in mechanical state, we measured the strain rate (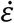) as a function of myosin concentration. In α-actinin-crosslinked networks, the strain rate exhibits a non-monotonic dependence on myosin concentration, reaching a maximum near R_myo_ ≈ 0.02 (Fig. 1l, SFig. 1). Thus, increasing motor abundance does not lead to a proportional increase in network deformation.

Interestingly, the non-monotonic dependence of strain rate on motor abundance suggests that the network does not respond to increasing motor activity as a passive material. Previous studies of active actomyosin systems have shown that contractility can depend non-monotonically on network connectivity (*C*) and rigidity (*G*), reflecting a balance between force transmission and mechanical resistance ^30,46^. At low connectivity, forces generated by motors are poorly transmitted through the network, whereas highly connected or rigid networks can resist deformation and suppress contractile motion. Similarly, our prior computational studies have demonstrated that monotonic changes in crosslinker mechanosensitivity and network connectivity can give rise to non-monotonic mechanical responses through mechanochemical feedback, emphasizing that active stress alone is insufficient to determine material behavior ^17^. Consequently, motor abundance alone does not uniquely determine the mechanical response of the system. Instead, the response depends on an evolving mechanical state, *X*. Here, *X* is not intended as a complete thermodynamic state variable, but rather as a coarse-grained description of the network properties most strongly implicated in force transmission, including effective connectivity (*C*), rigidity (*G*), and, more generally, the evolving architecture through which active stresses are generated and transmitted^47^.

If dissipation depends not only on motor abundance but also on the state of the receiving network, then energetic dissipation need not vary monotonically with motor abundance. Thus, the observed non-monotonic dissipation therefore emerges despite an approximately constant thermodynamic driving force, indicating that dissipation is regulated by the evolving mechanical state of the network rather than by Δ*μ*_*ATP*_ alone. This interpretation is consistent with our previous work showing that actin-network architecture can directly regulate the conversion of chemical energy into mechanical work ^14,15,25^. To test the possibility that system dissipation varies non-monotonically with motor abundance, we next directly measure heat dissipation using picowatt-resolution calorimetry.

### Crosslinker mechanics alter the relationship between motor abundance and active stress

We reconstitute a three-dimensional active actomyosin network by mixing purified G-actin, skeletal muscle myosin II, and crosslinkers such as α-actinin or fascin with ATP in F-buffer (Methods). Unlike the quasi-two-dimensional assay, the three-dimensional system does not exhibit large-scale contraction, reflecting differences in geometry and effective protein concentration (Fig. 2g, Supplementary Movie 2). The protein mixture is injected into a glass capillary mounted within a custom-built picowatt calorimeter. Heat dissipation, (Δ*Q*), is measured directly, with measurements acquired every 0.5 s, yielding the dissipated power 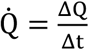. The calorimeter achieves a detection limit of (ΔQ = 100) pW^42^ (Fig. 2a, SFig. 2, Methods).

**Fig. 2.**
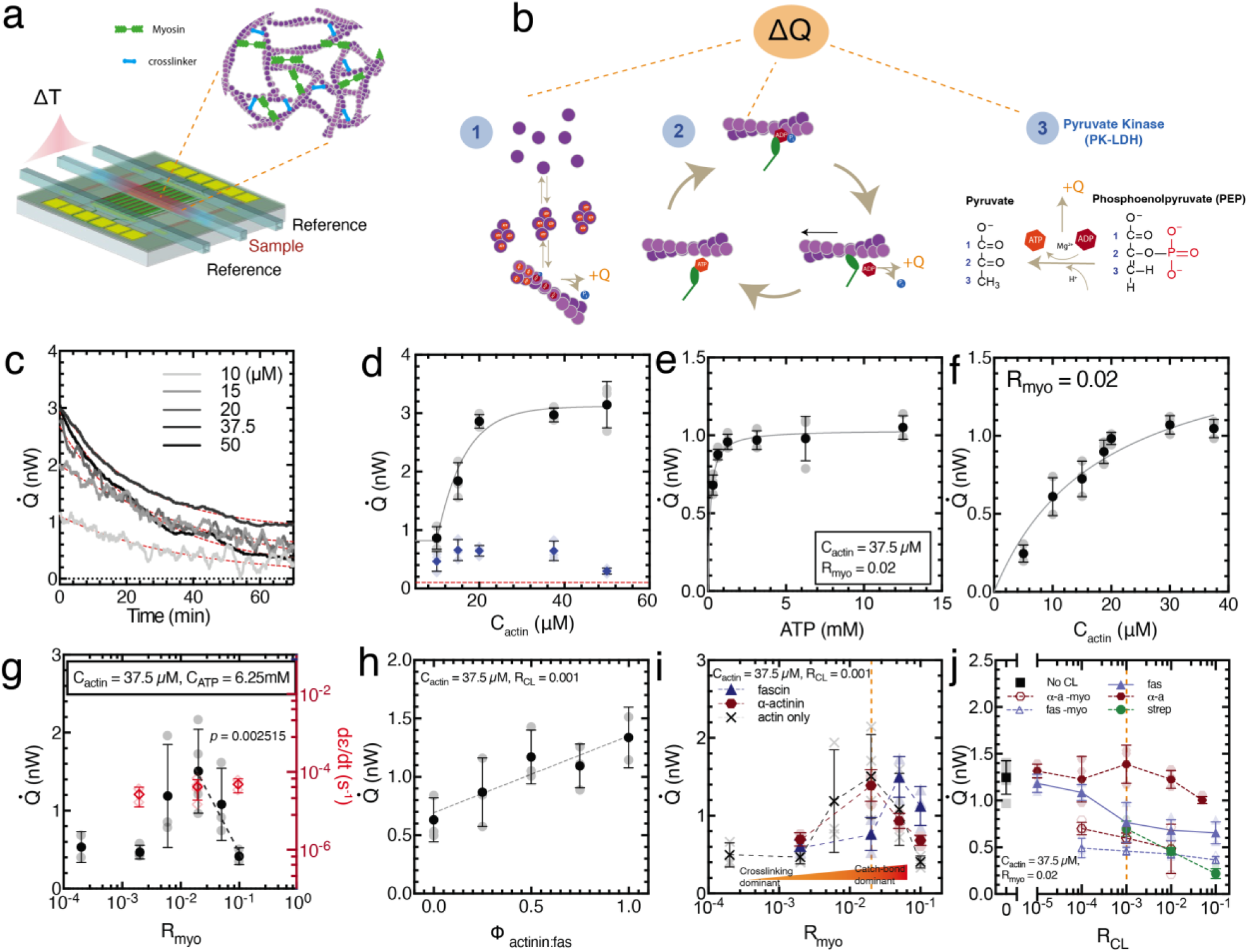
F-actin crosslinkers tune myosin-generated heat dissipation. (a) Schematic of calorimeter and the zoomed-in circle shows the actomyosin sample details in the capillary tube that contains the sample. (b) Schematic of ATPase activities from 1. actin polymerization; 2. myosin ATPase in the network; and 3. the chemical reaction of ATP regeneration system via Pyruvate assay. (c) Heat dissipation rate (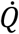) of actin polymerization at various actin concentrations over time. (d) Initial (black circles) and steady state (blue diamonds) heat dissipation rate (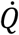) at various actin concentrations. Gray line is a Michaelis-Menten curve fit. N = 3 independent experiments for each condition. Error bars are the s.d. of the mean. (e) Heat dissipation rate of actomyosin network at various ATP concentrations (C_actin_ = 37.5µM, R_myo_ = 0.02.) Black line is the fit of Michaelis-Menten curve to data. N = 3 independent experiments for each condition. Error bars are the s.d. of the mean. (f) Heat dissipation rate of actomyosin network at various actin concentrations (R_myo_ = 0.02.) Black line is the fit of Michaelis-Menten curve to data. N = 3 independent experiments for each condition. Error bars are the s.d. of the mean. (g) Heat dissipation rate (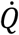) of actomyosin network at various myosin-to-actin ratios (C_actin_ = 37.5µM, C_ATP_ = 6.25mM.) N = 3 independent experiments for each condition. Error bars are the s.d. of the mean. *p* = 0.002515 for the downward trend using standard linear model least squares regression. (h) Heat dissipation rate (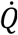) of actomyosin network under same crosslinking ratio R_CL_ = 0.001 but at different α-actinin:fascin ratios (Φ_*actin*:*fascin*_). Gray dashed line is a linear fit to the data. N = 3 independent experiments for each condition. Error bars are the s.d. of the mean. (i) Heat dissipation rate (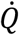) of actomyosin network under same crosslinking ratio R_CL_ = 0.001, C_ATP_ = 6.25mM, and C_actin_ = 37.5µM, at various R_myo_. N = 3 independent experiments for each condition. Error bars are the s.d. of the mean. (j) Heat dissipation rate (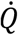) of actomyosin network at R_myo_ = 0.02, C_ATP_ = 6.25mM, and C_actin_ = 37.5µM, and various crosslinking ratio R_CL_. N = 3 independent experiments for each condition. Error bars are the s.d. of the mean.

The measured heat signal contains three principal contributions: (i) ATP hydrolysis associated with actin polymerization, (ii) ATP hydrolysis by myosin motor activity, and (iii) ATP regeneration through the pyruvate kinase/lactate dehydrogenase (PK-LDH) system and phosphoenolpyruvate (PEP) (Fig. 2b, SFig. 2, Methods). Upon actin polymerization alone, (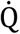) exhibits an exponential decay in time (Fig. 2c,d, Supplementary Materials). After characterizing the energetic contribution from actin polymerization, we introduce skeletal muscle myosin II dimers, which polymerize into thick filaments (Supplementary Materials).

We first establish the baseline response of the system by varying ATP concentration while maintaining constant actin and myosin concentrations. Under these conditions, ATP consumption follows the expected Michaelis-Menten kinetics, consistent with conventional enzyme behavior. This provides a useful reference because varying ATP concentration over a certain range changes the chemical driving without substantially altering the mechanical organization of the network. In contrast, varying motor abundance changes both the driving and the material itself:

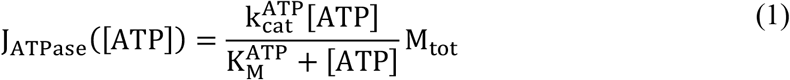

where 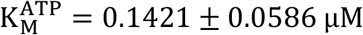. The resulting ATPase rates differ from previous measurements by approximately a factor of two ^48^ (Fig. 2e), which may arise from differences in assay conditions, including actin concentration and myosin isoform. Similarly, varying the actin concentration while maintaining constant myosin concentration yields Michaelis–Menten behavior,

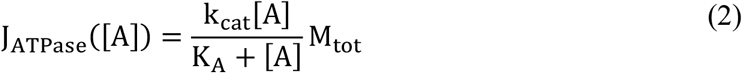

with K_A_ = 18.46 ± 6.73 μM, in agreement with previous studies ^49,50^(Fig. 2f).

To investigate the energetic consequences of motor activity, we next vary myosin concentration while maintaining a constant actin concentration, *R*_*myo*_. Under these conditions, the dissipated power exhibits a non-monotonic dependence on myosin concentration (Fig. 2g). At low motor density, dissipation increases with increasing myosin concentration, consistent with the addition of ATP-consuming motors to the network. Dissipation reaches a maximum near R_myo_ ≈ 0.02 and subsequently decreases at higher myosin concentrations. Thus, motor abundance and energetic dissipation are not simply proportional over the experimentally accessible range. Instead, the dissipation becomes influenced by system state:

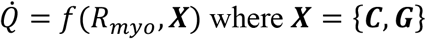

Separately, the dissipated power is proportional to the concentration of ATP-consuming myosin heads, (c_head_), the ATP turnover rate per motor, (*r*), the chamber volume (*V*), and the free energy released per ATP hydrolysis event, (ΔG), such that:

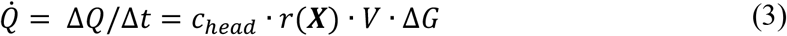

Using Eq. (3), we estimate an effective ATPase rate for the motor population (SFig. 3, Supplementary Materials). Because ATP turnover is known to depend on mechanical loading at the single-molecule level^51–54^, and because actin architecture can regulate energy conversion at the network level ^14^, ATPase rate provides information about the mechanochemical state of the motor population. However, it should not be interpreted as a direct quantitative measure of active stress. In dense actomyosin networks, the relationship between ATPase activity and active stress depends on motor occupancy, force sharing, network architecture, bundling, and remodeling dynamics.

When active stress is approximated as a motor force-dipole density, 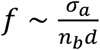, where *n*_*b*_ is the bound motor density and *d* is the force-dipole length, load-dependent motor kinetics suggest that ATPase activity can decrease with increasing motor load ^55–57^. Consequently, ATPase activity provides a qualitative indicator of motor mechanochemical state, but not a unique measure of active stress in concentrated active networks^14,43,58^.

The non-monotonic dependence of dissipation on myosin concentration suggests that increasing motor abundance alters not only the number of active motors, but also the mechanical state of the network in which those motors operate. To investigate how network mechanics may contribute to the observed energetic behavior, we next compare actomyosin networks crosslinked with either α-actinin or fascin using three-dimensional microscopy, rheology, and pico-calorimetry.

### State-dependent energy conversion produces non-monotonic dissipation

As noted above, the three-dimensional actomyosin networks exhibit minimal large-scale deformation over the range of motor concentrations examined, likely reflecting differences in geometry and effective protein concentration relative to the quasi-two-dimensional assay, as reported previously^23,29^ (Fig. 2g). Given the distinct mechanical responses observed for α-actinin- and fascin-crosslinked networks under active loading, we next quantify their energetic behavior using picowatt-resolution calorimetry^16^.

At fixed myosin concentration and total crosslinker density, increasing the fraction of α-actinin relative to fascin Φ_actinin:fascin_ results in a monotonic increase in heat dissipation (Fig. 2h). This observation indicates that catch-bond and slip-bond crosslinkers differentially regulate energy consumption despite being coupled to the same actomyosin motor system. We therefore ask whether crosslinker mechanics also influence how dissipation responds to increasing motor activity.

To address this question, we systematically vary myosin concentration while maintaining a constant crosslinker density. In both α-actinin- and fascin-crosslinked networks, the dissipated power, 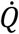, exhibits a non-monotonic dependence on motor concentration (Fig. 2i). Dissipation initially increases with motor abundance, reaches a maximum, and subsequently decreases at higher motor concentrations. However, the position of this maximum depends strongly on crosslinker identity, occurring at substantially higher myosin concentrations in fascin-crosslinked networks than in α-actinin-crosslinked networks. We interpret this maximum as marking the transition between two energetic regimes. Below the peak, increasing motor abundance primarily recruits additional ATP-consuming motors, resulting in approximately proportional increases in dissipation. Beyond the peak, further increases in motor abundance predominantly reorganize the mechanical state of the network, such that dissipation becomes increasingly governed by the evolving state (***X***) rather than by motor abundance alone. Thus, crosslinker mechanics do not simply shift the dissipation curve; they regulate the onset of state-dependent energy conversion by determining how much motor activity the network can accommodate before mechanochemical feedback begins to dominate.

The shift in the dissipation maximum raises the question of whether the observed behavior arises simply from differences in network connectivity or more fundamentally from the force-dependent mechanics of the crosslinkers themselves. To distinguish between these possibilities, we next examine how dissipation depends on crosslinker density. For fascin-crosslinked networks, heat dissipation decreases sharply near R_c_ ≈ 0.001, close to the expected bundling threshold and percolation critical point for three-dimensional actin networks^41,59^ (Fig. 2j). In contrast, α-actinin-crosslinked networks display only a modest decrease in dissipation as crosslinker density increases (Fig. 2j). Consequently, α-actinin-crosslinked networks dissipate more energy than fascin-crosslinked networks across nearly the entire range of crosslinking densities examined.

Together, these observations indicate that crosslinker mechanics substantially alter the energetic response of active actomyosin networks despite identical motor proteins and ATP availability. Consistent with this interpretation, the two network architectures exhibit different effective ATPase rates (SFig. 2). However, calorimetry alone cannot determine how these energetic differences arise from changes in the mechanical state of the network. To directly characterize the mechanical state of the networks, we next perform rheological measurements to quantify their viscoelastic response under motor activity (Fig. 3a).

**Fig. 3.**
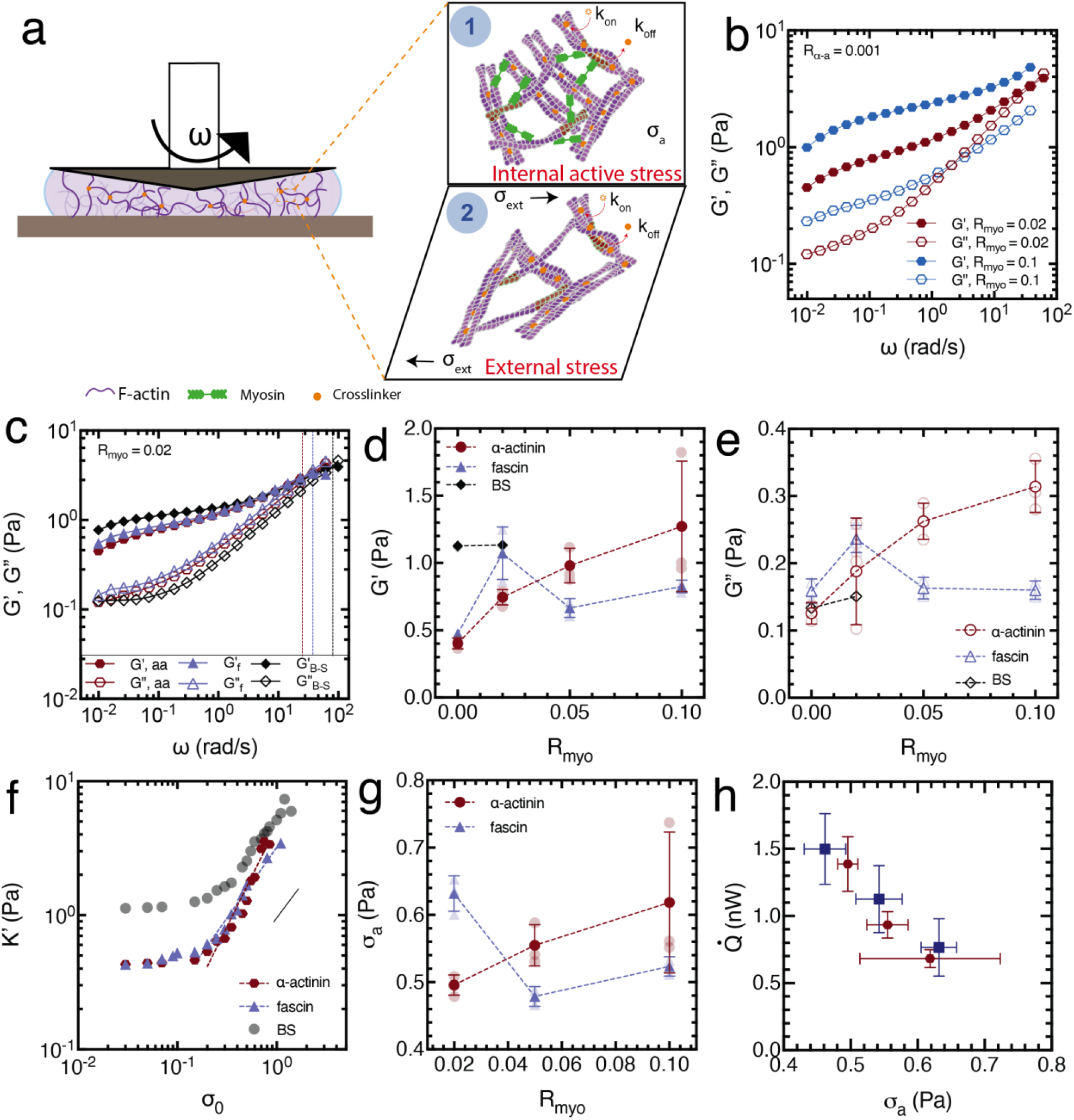
Crosslinker mechanics alter the relationship between motor abundance, active stress, and dissipation. (a) Schematic of crosslinked network inside a cone-and-plate rheometer for (1) under active stress (top right) and (2) under rheometer-induced prestress (bottom right). (b) Frequency sweep (*G*^′^, elastic modulus, and *G*^′′^, viscous modulus) of the α-actinin crosslinked network (R_CL_ = 0.001) under active stress induced by myosin (R_myo_ **=** 0.02, maroon, and 0.1, blue). (c) Frequency sweep (*G*^′^, elastic modulus, and *G*^′′^, viscous modulus) of the α-actinin (maroon hexagons), fascin (blue triangles), and biotin-streptavidin (black diamonds) crosslinked networks. Vertical dashed lines indicate the *G*^′^and *G*^′′^crossover frequency ω;_co_ for three cases. (d) Elastic modulus *G*^′^ of the three types of network with various R_myo_ at ω; = 0.01 rad/s. N = 3 independent experiments for each condition. (e) Viscous modulus *G*^′′^ of the three types of network with various R_myo_ at ω; = 0.01 rad/s. N = 3 independent experiments for each condition. (f) Differential elastic modulus K’ for the three types of networks as a function of prestress σ_0_. Black line is indicative of slope of 1. (g) Indicated active stress σ_a_ as a function of R_myo_ for the three types of networks. N = 3 independent experiments for each condition. (h) Total heat dissipation rate 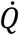 as a function of inferred active stress σ_a_. N = 3 for each condition. Error bars are the s.d. of the mean.

### Crosslinker mechanics alter the relationship between motor abundance and active stress

To characterize the material properties of crosslinked F-actin networks under active stress generated by myosin motors, we use a stress-controlled rheometer (Fig. 3a, Supplementary Materials). Previous studies have suggested that the mechanical response of actively stressed cytoskeletal networks is analogous to that of passive networks driven into the nonlinear regime through externally applied prestress^23,29,60^. Consistent with this picture, we observe that the viscoelastic moduli *G*’ and *G*’’ increase with increasing R_myo_ (Fig. 3b).

In linear frequency-sweep experiments, the crossover frequency ω_co_, defined by (*G*′ = *G*′′), provides a characteristic timescale separating solid-like and fluid-like behavior. At equivalent crosslinking densities, α-actinin-crosslinked networks exhibit significantly lower *ω*_*co*_ values than fascin-crosslinked networks, indicating more dynamic and adaptive mechanical behavior (Fig. 3c, SFig. 4). As a comparison, networks crosslinked through permanent biotin–streptavidin linkages display little or no crossover over the accessible frequency range and behave predominantly as elastic solids (Fig. 3c, SFig. 4).

As myosin density increases, both *G*′ and *G*′′ increase monotonically in α-actinin-crosslinked networks, consistent with stress-induced stiffening reported previously^23,60^ (Fig. 3d-e). In contrast, fascin-crosslinked networks exhibit a non-monotonic rheological response, with both moduli initially increasing and subsequently decreasing as motor concentration rises (Fig. 3d-e). These observations suggest that motor activity drives the two network architectures into mechanically distinct states.

To estimate the level of internally generated stress, we perform nonlinear rheology on passive networks of matched crosslinking density (Methods). We first apply a step shear prestress σ_0_ and allow the sample to reach a quasi-steady state. We then superimpose a small oscillatory stress perturbation δσ(t) = |δσ|e^iωt^, maintaining amplitudes within the linear response regime of the prestressed state |δσ| ≤ σ_0_/10^61–63^. The resulting strain response defines the complex differential modulus,

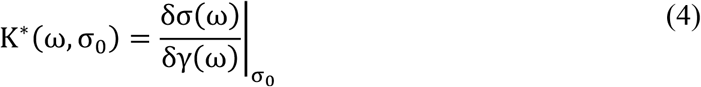

with K^′^(ω, σ_0_) = Re(K^*\**^) and K^′′^(ω, σ_0_) = Im(K^*\**^). In the limit of zero prestress, the differential modulus reduces to the conventional linear viscoelastic moduli, K^′^(ω, 0) = G^′^(ω)and K^′′^(ω, 0) = G^′′^(ω). The differential elastic modulus is fit using:

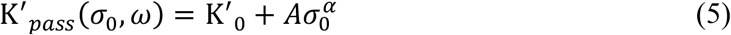

allowing the linear rheology of active networks to be mapped onto the stress–stiffening curves of passive networks. This procedure provides an estimate of the active stress associated with motor activity:

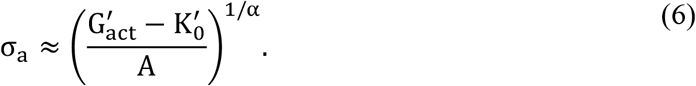

This mapping provides an effective active prestress that would produce the same linear elastic modulus in a passive network.

Using this analysis, we find that fascin-crosslinked networks exhibit larger inferred active stresses than α-actinin-crosslinked networks at low myosin concentrations (Fig. 3h). However, above R_myo_ ≈ 0.05, the inferred active stress in fascin networks decreases. Remarkably, this trend parallels the calorimetric measurements (Fig. 2i), where the energetic behavior of the two network types also diverges at elevated motor concentrations.

When dissipation is plotted against myosin concentration, the relationship is non-monotonic. In contrast, dissipation decreases approximately monotonically with inferred active stress (Fig. 3h). While myosin concentration serves as the experimentally imposed control parameter throughout this work, the active stress is an emergent quantity that depends on motor loading, network architecture, and mechanochemical feedback. Consequently, increasing motor abundance does not necessarily produce proportionally larger active stresses. Thus, the relevant causal sequence is not simply 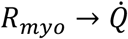, but rather 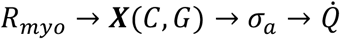, where *X* denotes the evolving mechanical state of the network, including connectivity *C*, rigidity *G*. The observed non-monotonicity therefore reflects the response of a reorganizing active material to increased motor abundance rather than a direct measurement of dissipation as a function of active stress alone. Rather, increasing motor abundance alters both the energetic and mechanical state of the network, indicating that dissipation depends on the coupled evolution of motor activity and network mechanics rather than on motor number alone.

Although calorimetry measures the total chemical energy dissipated, only a fraction is converted into measurable mechanical deformation. To quantify this partitioning, we estimate the mechanical power associated with network deformation. Thus, we use velocity fields obtained from three-dimensional PIV measurements to estimate the mechanical dissipation rate,

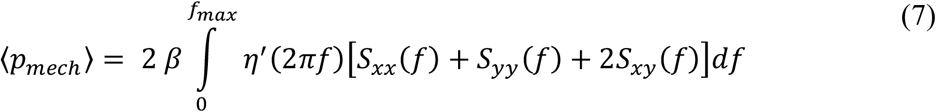

where β is the inverse thermal energy of the bath, *S*(*f*) is the power spectral density of the strain-rate field, and the dynamic viscosity is given by η^′^(2πf) = G^′′^(f)/(2πf) (Supplementary Materials). From this quantity we calculate the mechanical energy-conversion efficiency,

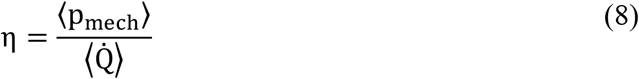

Mechanical energy-conversion efficiencies range from approximately 5 × 10^−5^ to 1.5 × 10^−4^, with fascin-crosslinked networks exhibiting higher efficiencies than α-actinin-crosslinked networks (SFig. 5). Thus, although both systems consume chemical energy through ATP hydrolysis, the fraction converted into measurable mechanical work depends strongly on the evolving network architecture and crosslinker mechanics.

### Distinct mechanical states exhibit similar dissipation

The preceding results suggest that motor abundance, active stress, and energetic dissipation become decoupled as the network reorganizes. We therefore asked whether mechanically distinct network states could nevertheless exhibit similar energetic costs. To further characterize the evolving mechanical state, we analyzed actin-network velocity correlations and estimated the entropy production rate (EPR) from the actin and myosin fluorescence channels using established nonequilibrium inference approaches^26,64,65^ (Methods). We use the EPR as an independent measure of nonequilibrium activity and as a lower bound on the total entropy production of the system, rather than as a complete measure of all dissipative processes. This interpretation assumes that the dominant contribution arises from myosin-driven ATP hydrolysis, while additional dissipative pathways, including filament turnover, network remodeling, and viscoelastic relaxation, contribute to the total dissipation measured by calorimetry (Fig. 2).

The estimated EPR exhibits a non-monotonic dependence on myosin concentration and displays similar values at low (R_myo_ = 0.002) and high (R_myo_ = 0.1) motor abundance (Fig. 4a,d). This trend parallels the behavior observed in the calorimetric measurements, suggesting that the inferred EPR captures a substantial component of the dominant dissipative dynamics. Because the EPR estimate is necessarily limited by the measured variables and underlying model assumptions, the observed degeneracy should be interpreted as a degeneracy in the inferred entropy production rather than as a demonstration that all microscopic dissipative pathways are identical.

Although these regimes exhibit comparable dissipation, they display distinct velocity-correlation length structures (*ℓ*_*v*_). At low motor abundance, velocity correlations remain relatively short-ranged, whereas at higher motor abundance the correlations extend over larger distances and are associated with increasingly coherent network flows (Fig. 4b,c). These observations indicate that mechanically distinct dynamical states *X* = (*C, G, ℓ*_*v*_) can nevertheless exhibit similar energetic costs (Fig. 4e).

**Fig. 4.**
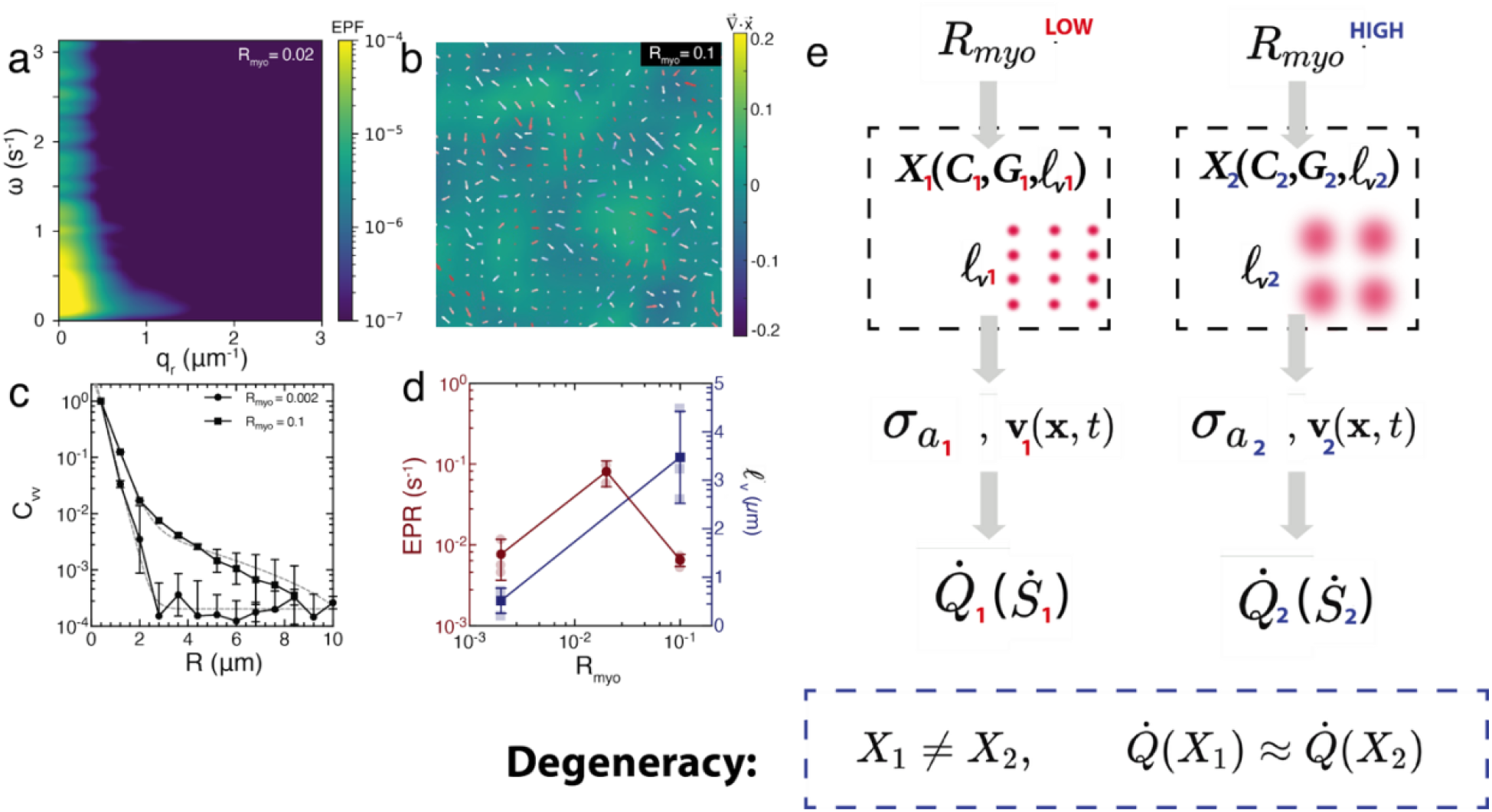
Degeneracy in dissipation accompanies distinct mechanical states. (a) Entropy production factor (EPF) for frequency (*ω*) and wave vector (*q*_*r*_), calculated from actin and myosin dynamics at intermediate motor concentration, R_myo_ = 0.02. The EPF is concentrated at low spatial frequencies, indicating that the dominant irreversible dynamics occur over large spatial scales. (b) Heatmap shows accumulative strain (colorbar) after 5 minutes of completion of polymerization. Quiver plot overlay shows the mean velocity within the 5 minutes. Arrow color indicates the degree of alignment of the velocity with its neighbors. Red-white-blue indicates ‘highly aligned’-‘not aligned’ – ‘anti-aligned’. R_myo_ = 0.1. (c) Azimuthally averaged velocity spatial correlation (*C*_*vv*_) as a function of radius *R*. N = 3 for both cases. Errorbars are the s.d. of the mean. Gray dashed lines are the exponential fit. (d) Entropy Production Rate (EPR, maroon, left y-axis) and characteristic length (*l*_*v*_, blue, right y-axis) as a function of R_myo_. N = 3 for all conditions. The EPR is the integrated value of the EPF. Errorbars are the s.d. of the mean. (e) Conceptual framework illustrating dissipation degeneracy. Different motor abundances generate distinct mechanical states (X = {*C, G, ℓ*_*v*_}characterized by different network connectivity (*C*), rigidity (*G*), and velocity-correlation length, *ℓ*_*v*_. These states produce different active stresses σ_a_ and velocity fields **v**(x, t), yet can exhibit comparable heat dissipation rates 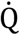 (and corresponding entropy-production estimates 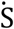). Thus, distinct mechanical states can map onto similar energetic costs, yielding a degeneracy between mechanical organization and dissipation in active actomyosin networks.

Although the present experiments do not directly resolve spatial stress fields, the combined rheological, velocity-correlation, morphological, and calorimetric measurements are consistent with a reorganization of force transmission as motor abundance increases. One interpretation consistent with the observed differences in velocity correlations, rheology, morphology, and dissipation is that force transmission becomes reorganized as motor abundance increases. Indeed, previous work showed that α-actinin-4 forms catch bonds whose lifetime increases under tension, allowing crosslinks to stabilize load-bearing regions and redistribute forces across actin networks^21,66^. In contrast, fascin promotes bundled architectures that undergo clustered contraction and coarsening ^41,45,67–70^, and a more localized mode of force transmission^68,69,71^. Within this framework, crosslinker-specific bond mechanics and network architecture provide one potential mechanism by which α-actinin-rich networks could support more distributed force transmission, whereas fascin-rich bundled networks could maintain more localized contractile stresses^72^.

These observations establish that energetic cost does not uniquely specify the mechanical state of an active actomyosin network. Instead, mechanically distinct states can converge to similar dissipation despite substantial differences in network organization and deformation dynamics. This dissipation degeneracy arises because motor abundance reorganizes the material itself, causing energetic cost to depend on the evolving mechanical state rather than actuator abundance alone.

## Discussion

The present study demonstrates that energy dissipation in active actomyosin networks depends non-monotonically on motor abundance and that mechanically distinct network states can exhibit similar energetic costs despite substantial differences in network organization, connectivity, deformation dynamics, and force transmission. Notably, while dissipation depends non-monotonically on myosin abundance, it varies approximately monotonically with the inferred active stress. This suggests that active stress functions as an emergent coarse-grained variable linking the evolving mechanical state of the network to its energetic cost.

Together, these findings reveal a degeneracy between energetic dissipation and mechanical state, indicating that energy consumption cannot be inferred from actuator abundance alone. Rather, motor abundance acts as a control parameter that reorganizes the material, altering the pathways through which forces are transmitted, and chemical energy is converted into work and heat. This conclusion extends previous observations that entropy production is maximized at intermediate motor abundance in quasi-two-dimensional actomyosin networks^26^ by demonstrating the same qualitative behavior using direct calorimetry and across distinct crosslinker architectures. Importantly, the non-monotonic energetic response is accompanied by systematic changes in rheological state, active stress, and velocity correlations, linking dissipation directly to the evolving mechanical organization of the network.

We interpret the maximum in dissipation as marking the onset of state-dependent mechanochemical feedback. Below this transition, increasing motor abundance primarily recruits additional ATP-consuming motors, producing larger active stresses and greater heat dissipation. Beyond the maximum, however, further increases in motor abundance predominantly reorganize the mechanical state of the network, altering force transmission and the mechanical loads experienced by individual motors. As a consequence, ATP turnover and dissipation decrease despite continued increases in motor abundance. Crosslinker mechanics therefore do not simply shift the dissipation curve; they regulate the motor abundance at which state-dependent energy conversion emerges.

One possible microscopic origin of this transition is that the force-dependent properties of the crosslinkers alter how motor-generated forces are transmitted through the network^73^. Previous studies have shown that *α*-actinin 4 forms catch bonds whose lifetime increases under tension, enabling load-bearing regions to remain mechanically connected under stress^21,66^. In contrast, fascin promotes bundled architectures that undergo clustered contraction and coarsening^68,69,71^. Within this framework, *α* −actinin-rich networks may distribute forces over larger regions of the network, whereas fascin-rich networks may support more localized force transmission^72^. Such differences in force organization could alter the mechanical load experienced by individual motors and thereby modify ATP turnover through load-dependent mechanochemical feedback.

More generally, our results suggest that motor abundance should be viewed as a control parameter rather than the thermodynamic driving force itself. The chemical potential difference associated with ATP hydrolysis remains approximately constant, whereas increasing myosin concentration reorganizes the receiving network and alters its connectivity, rigidity, and force-transmission pathways. Consequently, motor abundance, mechanical organization, active stress, and dissipation become distinct but coupled quantities whose relationships depend on the evolving state of the material. In this view, myosin abundance serves as an experimental control parameter, whereas active stress emerges from the coupled dynamics of motors and network architecture. The approximately monotonic relationship between active stress and dissipation therefore provides a physical explanation for why dissipation becomes non-monotonic when plotted against motor concentration.

This interpretation is consistent with our recent work demonstrating that F-actin architecture regulates the conversion of chemical energy into mechanical work ^14,74^ and with more recent studies showing that crosslinked networks regulate ATP consumption through load-dependent mechanochemical coupling ^15^. In those experiments, distinct crosslinkers produced substantially different ATP consumption rates despite identical motor proteins and ATP availability, indicating that network architecture can directly modulate energy conversion. Together, these findings support a framework in which motor abundance influences dissipation indirectly through the emergent mechanical state of the material. Here, *X* should not be interpreted as a complete thermodynamic state variable, but rather as a coarse-grained description of the network properties most relevant to force transmission and energy conversion. In the present system, these properties include network connectivity (*C*), rigidity (*G*), and ultimately the characteristic velocity-correlation length scale (*ℓ*_*v*_), although additional structural variables may also contribute (Fig. 4e).

From a thermodynamic perspective, the observed non-monotonic dissipation does not violate Onsager reciprocity or other near-equilibrium principles. Rather, it reflects the breakdown of assumptions that underlie linear irreversible thermodynamics, namely fixed transport coefficients and weak perturbations from equilibrium. In active materials, the mechanical state evolves together with motor activity, causing the effective coefficients governing force transmission and energy conversion to become state dependent. The experimentally imposed variable (motor abundance) therefore differs from the state variable governing energy conversion (active stress), because the latter emerges from the evolving mechanical organization of the network.

The resulting decoupling between motor abundance, mechanical organization, and energetic cost gives rise to what we term a *dissipation degeneracy*. The concept is analogous to degeneracy in biological systems, where structurally distinct configurations produce similar functional outputs^75^, but here the convergent observable is energetic dissipation rather than biological function. Specifically, mechanically distinct network states can exhibit comparable energetic dissipation, such that (X_1_ ≠ X_2_) while 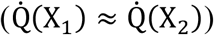 (Fig. 4e). In other words, energetic cost does not uniquely specify mechanical organization. States with distinct correlation lengths, rigidities, connectivities, and force-transmission pathways can converge to similar active stresses and therefore similar levels of dissipation despite differing substantially in their mechanical organization.

The distinction between motor abundance and active stress also provides an interesting contrast with classical muscle energetics, where ATP consumption, force generation, and heat production occur within a relatively fixed sarcomeric architecture ^76^. In contrast, active actomyosin networks continuously reorganize under motor loading ^22,69,77–81^, allowing motor abundance to modify the mechanical state of the material itself. Consequently, the relationship between motor activity and dissipation becomes state dependent rather than being determined solely by motor kinetics. Changes in force transmission similar to those observed here may therefore alter how chemical energy is partitioned between intracellular dissipation and external mechanical work^64^. Further, in non-muscle cells, forces generated by actomyosin are transmitted to adhesions, extracellular matrices, and neighboring cells^82^. Consequently, changes in force transmission similar to those observed here may also alter how chemical energy is partitioned between intracellular dissipation and external mechanical work, such as in the generation of traction stresses^64^. Understanding how network organization regulates this partitioning may therefore provide insight into migration, morphogenesis, and tissue mechanics.

## Acknowledgments

We are grateful for insightful discussions with Dr. Daniel Needleman. Z.G.S. acknowledges fundings and support from Yale PEB, his family, and friends during his exchange period at Harvard. Rheology experiments were carried out using the rheometer from Dr. Jing Yan’s lab at Yale University and paid for with Yale Startup Funds. This work was supported by funding ARO MURI W911NF-14-1-0403, the National Institutes of Health (NIH) R01 GR130179, Sloan Matter-to-Life G-2025-79182, and Human Frontiers Science Program (HFSP) grant number RGP012/2025 to M.P.M.

## Author Contributions

Z.G.S. & M.P.M designed and conceived the work. Z.G.S. & M.P.M. drafted the paper. Z.G.S, M.P.M, & J.J.V. edited the paper. Z.G.S. and A.P.T. performed experiments. Z.G.S. analyzed the data. J.Z. instructed on calorimeter operation. Z.G.S, M.P.M, J.J.V., & J.Z. participated in the discussion of the work. M.P.M. & J.J.V. supervised the work.

## Methods

### Preparation of small unilamellar vesicles (SUVs)

Lipids were combined by mixing 100 µL of egg phosphatidylcholine (Egg PC, 25 mg mL^−1^; Avanti Polar Lipids, 840051C) with 10 µL of 1,2-dihexadecanoyl-sn-glycero-3-phosphoethanolamine (DHPE, 0.5 mg mL^−1^), optionally conjugated to Oregon Green 488 (Invitrogen). All lipids were dissolved in chloroform and transferred to a clean glass vial pre-equilibrated with argon to minimize oxidative degradation. The solvent was removed under a gentle stream of argon, yielding a uniform lipid film coating the bottom of the vial. The dried lipid film was rehydrated with 5 mL of vesicle buffer (100 mM NaCl, 20 mM HEPES, pH 7.5) and vortexed until the suspension became turbid. The dispersion was then sonicated in a bath sonicator for approximately 1 h, or until the solution turned optically clear, indicating the formation of small unilamellar vesicles suitable for bilayer formation.

### Fluorescent labeling of skeletal muscle myosin

Skeletal muscle myosin (Heavy Meromyosin from rabbit, Cytoskeleton Inc.) is fluorescently labeled with Alexa Fluor 647 C2 Maleimide under reducing conditions. Initially, myosin is reduced in a labeling buffer containing 50 mM HEPES, 0.5 M KCl, 1 mM EDTA, and 10 mM DTT at pH 7.6. Following reduction, the sample is dialyzed overnight against the same buffer without DTT. After dialysis, the solution is centrifuged to eliminate any insoluble components. The resulting supernatant is reacted with Alexa Fluor C2 Maleimide at a 5:1 molar ratio of dye to myosin. Labeling is carried out at 4°C for one hour, after which the reaction is quenched by adding 1 mM DTT. The labeled protein is purified using a desalting column (Pierce, 5K MWCO, 5 mL). Absorbance readings at 280 nm and 647 nm are then used to calculate the degree of labeling. This protocol is adapted from Verkohovsky and Borisy^1^.

### 2D actomyosin contraction experiments

Each chamber is of cylindrical shape, with diameter of 12mm. The top and bottom piece are magnetically locked, with a rubber piece sealing the middle and a glass slide sandwiched in between. The glass slide is first washed with 50% ethanol to clean any residues left on the surface. The slide is then exposed under UV light for 5 min to induce hydrophilicity to the surface. 300 μL of SUV solution is added to the chamber. Once the surface of the glass is coated with lipid bilayer, we take 100 μL solution out and wash the chamber with 400 μL of 1x F-buffer (10 mM imidazole, 1 mM MgCl_2_, 50 mM KCl, 2 mM EGTA, 0.5 mM ATP, pH = 7.5). Dark G-actin (Cytoskeleton) is mixed with rhodamine labelled F-actin (20% fluorescent, Cytoskeleton) to a final molar concentration of 1.4 µM, and is stabilized with 1 μM phalloidin (Cytoskeleton) and crowded to the surface of a 97% Egg Phosphatidyl Choline (Avanti Polar Lipids)/3% FITC-DHPE (Molecular Probes) phospholipid bilayer, using 0.2% 14,000 MW methyl-cellulose (Sigma, 15 cP) as a depletion agent (Fig. 1A). The actin mixed soup is placed in an Eppendorf tube on ice for 1 hour to reach full polymerization (50 μL protein mix in total). Once F-actin is polymerized, it is added to the chamber, along with methyl-cellulose. Once the F-actin network is crowded onto the surface, we add crosslinkers *α*-actinin or fascin of various concentrations. Then, Skeletal muscle myosin II (various concentrations), labelled with Alexa Flour 647nm C2 Maleimide (Molecular Probes) is added in solution in dimeric form, which polymerizes into thick filaments onto the F-actin. This results in a two-dimensional actomyosin network.

### Removing non-catalytically active skeletal muscle myosin

To isolate only catalytically active myosin dimers for experimental use, myosin is subjected to a selective centrifugation process in the presence of polymerized actin. First, actin is polymerized for one hour at 4°C in a high-salt environment (1X F-buffer supplemented with 4 M KCl) and stabilized using phalloidin. After polymerization, ATP is added to reach a final concentration of 1 mM, followed by the addition of freshly thawed myosin. This actin-myosin mixture is incubated at 4°C for 10 minutes, then centrifuged at 128,360 × g for 30 minutes. During centrifugation, enzymatically inactive myosin remains bound to the F-actin network and sediments, while active myosin motors detach and remain in the supernatant, which is then collected. The concentration of active myosin in the supernatant is quantified by measuring absorbance at 647 nm, based on a pre-determined labeling efficiency. Myosin is freshly prepared for each experiment and used within 24 hours.

### Microscopy

The image stack data are collected using Leica DMi8 inverted microscope equipped with a 63×, 40×, or 20×, and 1.4, 1.3, and 0.75-NA oil immersion lens respectively (Leica Microsystems), a spinning-disk confocal (CSU22; Yokagawa), and sCMOS camera (Zyla; Andor Technology) controlled by Andor iQ3 (Andor Technology). Image time series stack data are taken with time interval of 0.5-10 seconds.

### Pico-calorimetry Measurements

Metabolic heat was quantified using a custom micromachined capillary-based picocalorimetry system, as previously described^42^ (Zheng et al., *PNAS*, 2026). Three borosilicate glass capillaries (400 µm × 400 µm outer dimension, 100 µm wall thickness) were bonded onto gold-coated regions of a silicon-nitride membrane integrating two Nichrome/Constantan thermopiles and a tungsten micro-heater. One capillary was loaded with the biological sample, and two identical capillaries containing pure water served as thermal references. Heat generated by the sample produced a temperature differential relative to the references, which was converted into a voltage by the thermopiles through the Seebeck effect. The monitored sensing volume within the sample capillary was ~80 nL.

The calorimeter assembly was mounted on a custom PCB and placed inside a thermally insulated vacuum chamber evacuated to ~10^−5^ Torr to suppress convective heat loss and reduce electrical noise. Measurements were performed at 25 °C. Thermopile voltages were acquired using two Keithley 2182A nanovoltmeters operated at 6 PLC with power-line synchronization and auto-zero enabled. Data were sampled at 2.31 Hz and processed using a moving-average filter. Under these measurement conditions, the system exhibited an effective noise floor corresponding to a power sensitivity of ~85 pW.

### Experimental setup and procedures for rheology experiments

Rheological measurements of actin networks were performed using a stress-controlled rheometer (Anton Paar M502). Actin was first prepared following the composition detailed in Table 1, and the final sample mixture (Table 2) was assembled to a total volume of 1000 µL. After gentle mixing, the solution was loaded onto the preheated bottom plate maintained at 25 °C. Both the cone and plate surfaces were sandblasted to prevent sample slippage. Following equilibration, several drops of silicone oil were added around the sample edge to minimize evaporation, and a solvent trap was placed over the geometry for additional protection. Gelation kinetics were monitored under the parameters described in the main text. After gelation reached a steady plateau in both elastic (*G*′) and viscous (*G*″) moduli (typically after ~2 h), measurements in the linear and nonlinear regimes were performed. After each experiment, the setup was thoroughly cleaned with sequential rinses of 70% ethanol, water, ethanol, and water to ensure complete removal of residual protein and prevent cross-contamination between runs.

### Experimental setup and procedures for microscopy experiments

Unlabeled rabbit skeletal muscle actin (>99% purity; Cytoskeleton, Inc.)—hereafter referred to as dark actin—was reconstituted in 1× G-buffer (see Table 3) to a concentration of 10 mg mL^−1^. Rhodamine-labeled actin (Cytoskeleton, Inc.) was prepared using the same procedure. Both preparations were depolymerized for over 24 h at 4 °C in the dark (see Table 3). The two actin species were then mixed at a 9:1 ratio (dark:rhodamine) to yield G-actin. Polymerization was initiated by combining the protein mixture with polymerization buffer (Table 4). Glucose oxidase, catalase (GOC), and glucose were added as an oxygen scavenging system to minimize photobleaching during imaging.

Samples were loaded into a custom four-well round chamber, each well comprising a cylindrical cavity sealed by a 12 mm glass coverslip and a rubber spacer to prevent leakage. Coverslips were sequentially cleaned with 70% ethanol, dried, and exposed to UV light for 5 min to render the surface hydrophilic. To reduce actin adsorption, surfaces were coated with small unilamellar vesicles (SUVs; see **Preparation of small unilamellar vesicles (SUVs)** in the **Methods** subsection). After gentle mixing (30 s), 200 µL of the protein mixture was injected into each well. Samples were imaged using a Leica confocal microscope (see *Microscopy* section).

### ATP-regeneration system

ATP regeneration system was used in all experiments, containing ATP at the indicated concentrations (0.625–12.5 mM), 40 mM phosphoenolpyruvate (PEP), and a coupled pyruvate kinase/lactate dehydrogenase (PK/LDH) enzyme mixture (Sigma-Aldrich). The PK/LDH enzymes were mixed at a 1:1 ratio (1 mg mL^−1^ stock concentration), and 2.5 µL of the enzyme mixture was added per reaction volume of 800 µL.

### Particle Image Velocimetry (PIV)

Particle image velocimetry (PIV) is applied on the fluorescent actin images in MATLAB (mPIV, https://www.mn.uio.no/math/english/people/aca/jks/matpiv/). The extent of contraction is calculated by defining a mean strain:

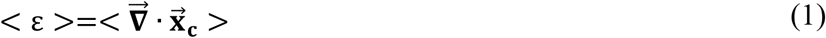

as the divergence of the displacement field. The data is analyzed with PIV window size 32 and overlap 0.5. Window size of 16 and 64 have also been used to generate the data, and eventually determined that 32 is the best parameter value because of the high signal-to-noise ratio.

### 2D image thresholding method for calculating bundling parameter

For each microscopy image of F-actin network, a line scan of approximately 20 μm is done to the image and the fluorescence intensity is extracted from the line scan. The bundling parameter λ is calculated using the equation:

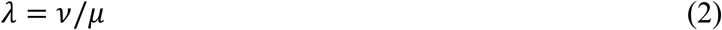

in which *v* is the standard deviation of the fluorescence intensity of the line scan, and *μ* the mean of the fluorescence intensity of the same line scan.

### Autocorrelation analysis

After the F-actin network reached a steady state, time-lapse fluorescence image sequences were acquired with randomly chosen starting times. To quantify the temporal dynamics of the network at the global level, we computed the traditional temporal autocorrelation of the fluorescence intensity using custom MATLAB scripts.

For each frame, the fluorescence intensity was spatially averaged over all pixels in the image, yielding a single intensity time series *I*(*t*). The mean intensity ⟨*I*⟩was then subtracted to isolate temporal fluctuations, δ*I*(*t*) = *I*(*t*) − ⟨*I*⟩. The normalized temporal autocorrelation function was calculated according to

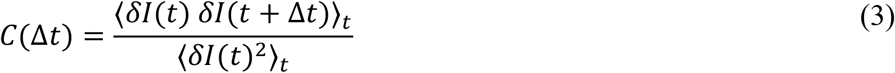

where ⟨⋅⟩_*t*_denotes an average over all valid time points separated by a delay Δ*t*. This normalization ensures *C*(0) = 1. Because the number of statistically independent pairs decreases with increasing delay time, the autocorrelation at large Δ*t* is increasingly affected by finite-sampling noise and was not interpreted quantitatively.

The experimentally measured autocorrelation functions were fit using a double-exponential decay model,

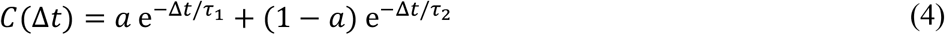

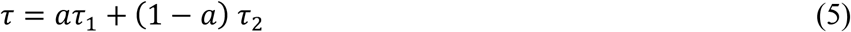

where *a* and 1 − *a* are the relative amplitudes of the two relaxation modes, and *τ*_1_and *τ*_2_are their associated characteristic times. Fitting was restricted to the initial portion of the correlation curve (typically the first 30% of the available delay times), as long-lag data points are dominated by statistical uncertainty arising from finite acquisition length.

### Mechanical power calculation using PIV velocity field

Defining 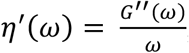, this is:

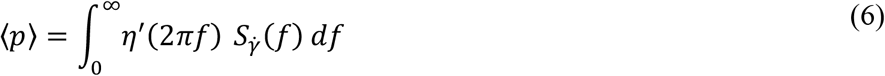

For a 2D/3D velocity field, using the strain-rate tensor 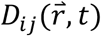 and its PSDs:

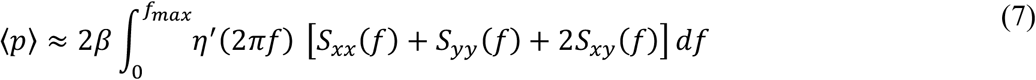

For more details, please refer to Supplemental Materials.

### Velocity spatial correlation analysis

Equal-time spatial velocity correlations were computed from particle image velocimetry (PIV) data using a custom MATLAB pipeline. The velocity correlation function was defined as

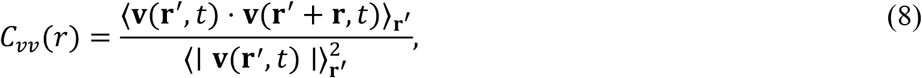

where **v**(**r**^′^, *t*)denotes the local velocity fluctuation at position **r**^′^and time *t*, and ⟨⋅⟩_**r**′_ indicates spatial averaging over all interrogation windows.

Velocity fields in the *x* and *y* directions (*v*_*x*_, *v*_*y*_) were first preprocessed to handle missing values arising from PIV failures. NaN entries were replaced using MATLAB’s fillmissing function with linear interpolation to ensure spatial continuity of the velocity field.

To compute the correlation efficiently and without directional bias, the calculation was performed in Fourier space. For each time point, the mean velocity was subtracted from the raw velocity field **v**_0_(**r**^′^, *t*)to obtain velocity fluctuations,

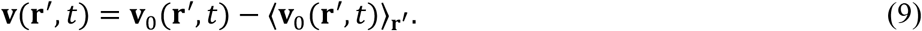

The resulting field was then normalized by its spatial root-mean-square magnitude,

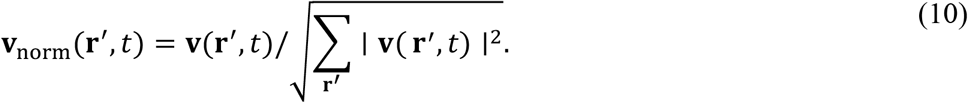

A two-dimensional fast Fourier transform (FFT) was applied to the normalized velocity field, and the autocorrelation was obtained via inverse FFT of the power spectrum:

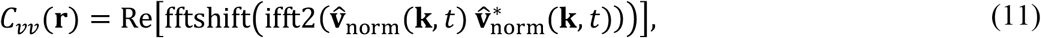

where 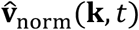 denotes the Fourier-transformed velocity field and *\** indicates complex conjugation. The fftshift operation was used to center the zero-frequency component.

The resulting two-dimensional correlation map was radially averaged using a custom radialavg routine to obtain *C*_*vv*_(*r*). The decay of the correlation function was quantified by fitting to a single-exponential form,

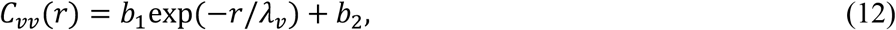

where *λ*_*v*_ defines the characteristic velocity correlation length.

### Entropy production calculation

The entropy production factor (EPF) and entropy production rate (EPR) are:

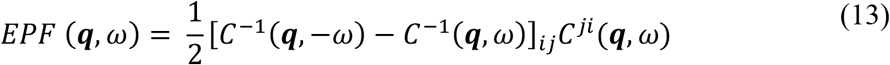

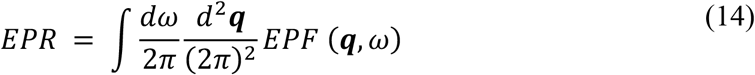

where *C*^*ij*^(***q***, *ω*) is the dynamic structure factor (for more derivations, please refer to Seara, Machta, and Murrell, 2021). The entropy calculation and analyses are carried out using customized Python script utilizing *frequent* and *freqentn* package developed by Dr. Daniel S. Seara (https://github.com/lab-of-living-matter/freqent/tree/epf_paper/freqent).

### Statistical analysis

Unless otherwise stated in the figure caption, pairwise comparisons were performed using Welch’s test, and linear fits were performed using standard linear model with least squares regression. Data displayed as a single value with an error bar are presented as mean ± standard deviation. Fitted lines are shown where applicable, with statistical significance indicated by the quoted p value. The symbols *, **, and *** in the figures denote p < 0.05, 0.01, and 0.001, respectively.

## Data availability

Source data are provided with this paper. Raw data supporting the findings of this manuscript are available from the corresponding authors upon reasonable request. A reporting summary for this Article is available as a Supplementary Information file.

## Code availability

Code supporting the findings of this manuscript are available from the corresponding authors upon reasonable request. A reporting summary for this Article is available as a Supplementary Information file.

