## Supplemental Materials for "State-dependent energy conversion produces degenerate dissipation in active actomyosin networks"

### Supplementary Materials for : **State-dependent energy conversion produces degenerate dissipation in active actomyosin networks**

Zachary Gao Sun[1,2,3,4], Juanjuan Zheng[4], A. Pasha Tabatabai[2,5], Joost J. Vlassak[4], and Michael Murrell[1,2,3,5,*]

[1] Department of Physics, Yale University, 217 Prospect Street, New Haven, Connecticut 06511, USA

[2] Systems Biology Institute, Yale University, 850 West Campus Drive, West Haven, Connecticut, 06516, USA

[3] Integrated Graduate Program in Physical and Engineering Biology, Yale University, New Haven, Connecticut 06520, USA

[4] John A. Paulson School of Engineering and Applied Sciences, Harvard University, Cambridge, MA, USA

[5] Department of Biomedical Engineering, Yale University, 55 Prospect Street, New Haven, Connecticut 06511, USA

**Experimental Details**

**Microscopy and rheology experiment buffer conditions**

**Rheology experiment:**

Table 1.

Actin overnight depolymerization:

| | Skeletal muscle actin acetone powder | ATP | Tris HCl | DTT | $NaN_3$ | $CaCl_2$ | pH |
|---|---|---|---|---|---|---|---|
| Final conc. | 10 mg/mL (238μM) | 0.2 mM | 2 mM | 0.2 mM | 0.005% | 0.2 mM | 7.0 |

Table 2.

Actin polymerization on rheometer:

| | Skeletal muscle actin | ATP | Tris HCl | $MgCl_2$ | KCl | $CaCl_2$ | crosslinker |
|---|---|---|---|---|---|---|---|
| Final conc. | 1.5625 mg/mL (37.5 μM) | 6.25 mM | 2 mM | 2 mM | 100 mM | 0.2 mM | $R_c * C_{actin}$ |

**Microscopy experiment:**

Table 3.

Actin overnight depolymerization:

| | Skeletal muscle actin acetone powder + rhodamine labeled actin (total 15% labeled) | ATP | Tris HCl | DTT | $NaN_3$ | $CaCl_2$ | pH |
|---|---|---|---|---|---|---|---|
| Final conc. | 10 mg/mL (238μM) | 0.2 mM | 2 mM | 0.2 mM | 0.005% | 0.2 mM | 7.0 |

Table 4.

Actin polymerization in chamber:

| | Skeletal muscle actin (15% rhodamine labeled) | ATP | Tris HCl | $MgCl_2$ | KCl | 50x Glucose Oxidase & Catalase* | 50x Glucose* | $CaCl_2$ | crosslinker |
|---|---|---|---|---|---|---|---|---|---|

| | | | | | | | | | |
|---|---|---|---|---|---|---|---|---|---|
| Final conc. | 1.5625 mg/mL (37.5 μM) | 0.5 mM | 2 mM | 2 mM | 100 mM | 6 μL | 6 μL | 0.2 mM | $R_c * C_{actin}$ |

*Note:

1. 50x Glucose Oxidase & Catalase is a mixture of Glucose Oxidase (CALBIOCHEM) of 5mg/mL with Catalase (CALBIOCHEM) of 0.9mg/mL[1,2]. (0.1 mg/mL and 0.018 mg/mL in experimental chamber respectively).

2. 50x Glucose contains 5 mg/mL of D-glucose mixed with 0.5% 2-mercaptoethanol (βME)[1,2]. (0.1mg/mL in experimental chamber).

**Equations & Calculations**

Catch bond & Slip bond binding rate:

The binding kinetics of the two types of bonds can be described as follow:

For slip bond:

$$k_{off}(f) = \frac{1}{\tau_{c0}} e^{\frac{f \Delta x_c}{k_B T}} \quad (1)$$

, and for catch bond:

$$k_{off}(f) = \frac{1}{\tau_{c0}} e^{-\frac{f \Delta x_c}{k_B T}} \quad (2)$$

.

**Kinetic equations**

Actin polymerization

Actin polymerization follows the following kinetic equation[3]:

$$A_m \cdot ATP + end \rightleftharpoons A_F \cdot ATP(end) \rightleftharpoons A_F \cdot ADP \cdot P_i + \Delta Q \quad (3)$$

, in which $A_m$ is G-actin, $A_F$ F-actin, $end$ filament end, and $P_i$ phosphate released from $ATP$ hydrolysis.

Myosin ATP hydrolysis:

$$A \cdot M + ATP \rightharpoonup A + M \cdot ATP \quad (4)$$

$$M \cdot ATP \rightleftharpoons M \cdot ADP \cdot P_i + A \rightleftharpoons A \cdot M \cdot ADP \cdot P_i \rightharpoonup A \cdot M \cdot ADP + P_i + \Delta Q \quad (5)$$

, in which $A$ is actin, $M$ myosin. Eqn. (4) shows myosin detachment from actin through ATP binding, and eqn. (5) shows the myosin power stroke and heat dissipation $\Delta Q$ during $P_i$ release, where chemical free energy is converted into mechanical work or stress and dissipated as heat.

ATP regeneration system (PH-LDH +PEP).

$$ATP + H_2O \rightharpoonup ADP + P_i \tag{6}$$

$$ADP + PEP \rightharpoonup ATP + pyruvate \tag{7}$$

, which leaves the net reaction to be:

$$PEP + H_2O \rightharpoonup pyruvate + P_i \tag{8}$$

Chamber volume V = 80 nL.

ATP regeneration via pyruvate kinase (PK-LDH).

Net reaction per ATP turnover: PEP + $H_2O$ → pyruvate + $P_i$.

Use |ΔG_net| ≈ 60 $kJ \cdot mol^{-1}$.

Heat dissipation rate during the process:

$$\dot{Q} = c_{head} \cdot r \cdot V \cdot 10^{-6} \cdot \Delta G$$

, with units:

c_head [μM] · r [$s^{-1}$] · V [L] · ΔG [$J \cdot mol^{-1}$] × ($10^{-6}$ $mol \cdot L^{-1}$ per μM).

Converting volume to nL and power to nW yields:

$$\dot{Q}[nW] = c_{head} \cdot r \cdot V_{nL} \cdot \Delta G_{kJ} \ / \ 1000$$

For $V_{nL}$ = 80 and $\Delta G_{kJ}$ = 60:

$$coefficient = \frac{(80 \times 60)}{1000} = 4.8$$

**Per turnover:**

Per ATP:

$q = \frac{\Delta G}{N_A} \approx 60{,}000\ J \cdot mol^{-1} / 6.022 \times 10^{23} \approx 1.03 \times 10^{-19}$ J.

Table A — Dissipation and inferred $c_{head} \cdot r$ ($\Delta G$ = 60 $kJ \cdot mol^{-1}$)

| myosin:actin ($R_{myo}$) | Dissipation P (nW) | c_head × r ($\mu M \cdot s^{-1}$) |
|---|---|---|
| 0.0002 | 0.53 | 0.110 |

| | | |
|---|---|---|
| 0.002 | 0.47 | 0.098 |
| 0.006 | 1.19 | 0.248 |
| 0.02 | 1.50 | 0.312 |
| 0.05 | 1.08 | 0.225 |
| 0.1 | 0.42 | 0.087 |

At constant myosin concentration ($R_{myo} = 0.02$):

1) Actin only

| Condition | $\dot{\boldsymbol{Q}}$ (nW) | c_head × r (μM·s⁻¹) |
|---|---|---|
| Actin only | 1.2469 | 0.2598 |

2) Fascin-crosslinked network

| Fascin:Actin | $\dot{\boldsymbol{Q}}$ (nW) | c_head × r (μM·s⁻¹) |
|---|---|---|
| 1e-05 | 1.1823 | 0.2463 |
| 0.0001 | 1.0904 | 0.2272 |
| 0.001 | 0.7666 | 0.1597 |
| 0.01 | 0.7089 | 0.1477 |
| 0.1 | 0.6551 | 0.1365 |

3) α-Actinin–crosslinked network

| α-Actinin:Actin | $\dot{\boldsymbol{Q}}$ (nW) | c_head × r (μM·s⁻¹) |
|---|---|---|
| 1e-05 | 1.3147 | 0.2739 |
| 0.0001 | 1.2281 | 0.2559 |
| 0.001 | 1.3377 | 0.2787 |
| 0.01 | 1.2243 | 0.2551 |
| 0.05 | 1.0158 | 0.2116 |

**Mechanical power calculation using PIV velocity field derived from visco-elastic dissipation**

1) Linear viscoelastic stress–strain relation

In shear, the stress is related to strain history by Boltzmann superposition:

$$\sigma(t) = \int_{-\infty}^{t} G(t-\tau)\ \dot{\gamma}(\tau)\, d$$

In frequency space:

$$\sigma(\omega) = G^{*}(\omega)\, \gamma(\omega), \quad G^{*} = G' + iG''$$

For sinusoidal strain $\gamma(t)=\gamma_0 \sin(\omega t)$, one finds:

$$\langle p_{mech} \rangle = \frac{1}{2}\, \omega\, \gamma_0^2\, G''(\omega)$$

Since

$$\Gamma(\omega) = \int_{-\infty}^{\infty} \gamma(t) e^{-i\omega t} dt$$

Parseval/Wiener-Khintchine theorem gives:

$$\frac{1}{2\pi T} |\Gamma(\omega)|^2 = S_{\gamma}^{(2)}(\omega)$$

Here $S_{\gamma}^{(2)}(\omega)$ is the two-sided PSD, and $T$ the averaging window.

Since $f = \frac{\omega}{2\pi}$,

$$S_{\gamma}(f) = 2 S_{\gamma}^{(2)}(2\pi f)$$

, and

$$< \gamma^2 > = \int_{0}^{\infty} S_{\gamma}(f)\, df$$

.

Using $S_{\gamma}(f)$ for one-sided PSD ($f \geq 0$, $\omega = 2\pi f$):

$$\langle p_{mech} \rangle = \int_{0}^{\infty} [\, 2\pi f\, G''(2\pi f)\, ]\; S_{\gamma}(f)\, df$$

Since $S_{\dot{\gamma}}(f) = (2\pi\, f)^2 S_{\gamma}(f)$:

$$\langle p_{mech} \rangle = \int_{0}^{\infty} \frac{G''(2\pi f)}{2\pi f}\, S_{\dot{\gamma}}(f)\, df$$

Defining $\eta'(\omega) = \frac{G''(\omega)}{\omega}$, this is:

$$\langle p_{mech} \rangle = \int_0^{\infty} \eta'(2\pi f)\; S_{\dot{\gamma}}(f)\, df$$

For a 2D/3D velocity field, using the strain-rate tensor $D_{ij}(\vec{r}, t)$ and its PSDs:

$$\langle p_{mech} \rangle \approx 2\beta \int_0^{f_{max}} \eta'(2\pi f)\; \left[S_{xx}(f) + S_{yy}(f) + 2S_{xy}(f)\right] df$$

**Derivation from $\boldsymbol{v(\vec{r}, t)}$ to strain-rate tensor $\boldsymbol{D_{ij}(\vec{r}, t)}$ to PSD $\boldsymbol{S_{xx}(f)}$.**

Strain-rate tensor from $v(\vec{r}, t)$:

$$D_{ij}(\vec{r}, t) \;=\; \frac{1}{2}\left(\partial_i\, v_j(\vec{r}, t) \;+\; \partial_j\, v_i(\vec{r}, t)\right)$$

Key components:

$$D_{xx} \;=\; \partial_x\, v_x,\;\; D_{yy} \;=\; \partial_y\, v_y,\;\; D_{xy} \;=\; \frac{1}{2}\left(\partial_x\, v_y \;+\; \partial_y\, v_x\right)$$

Temporal Fourier transforms at a pixel (two-sided, angular frequency ω):

$$V_j(\vec{r}, \omega) \;=\; \int_{-\infty}^{\infty} v_j(\vec{r}, t) e^{-i\omega\, t}\, dt$$

$$D_{ij}(\vec{r}, \omega) \;=\; \int_{-\infty}^{\infty} D_{ij}(\vec{r}, t)\; e^{-i\omega\, t}\, dt$$

Spatial derivatives commute with the temporal FT:

$$D_{ij}(\vec{r}, \omega) \;=\; \frac{1}{2}\left(\partial_i\, V_j(\vec{r}, \omega) \;+\; \partial_j\, V_i(\vec{r}, \omega)\right)$$

For the xx component:

$$D_{xx}(\vec{r}, \omega) \;=\; \partial_x\, V_x(\vec{r}, \omega)$$

Two-sided temporal PSD at a pixel (window T):

$${S_{xx}}^2(\vec{r}, \omega) \;=\; \frac{1}{2\pi\, T}\; |D_{xx}(\vec{r}, \omega)|^2$$

Parseval consistency (time average):

$$\left\langle {D_{xx}}^2(\vec{r},t) \right\rangle_t = \frac{1}{2\pi} \int_{-\infty}^{\infty} {S_{xx}}^2(\vec{r},\omega)\, d\omega$$

One-sided PSD in Hz ($f \geq 0$, $\omega = 2\pi f$), with spatial averaging over $\Omega$:

$$S_{D_{xx}}(\vec{r},f) = 2\, {S_{D_{xx}}}^2(\vec{r},2\pi f)$$

$$S_{xx}(f) = \left\langle S_{D_{xx}}(\vec{r},f) \right\rangle_{\boldsymbol{x}\in\Omega}$$

#### Supplementary figures

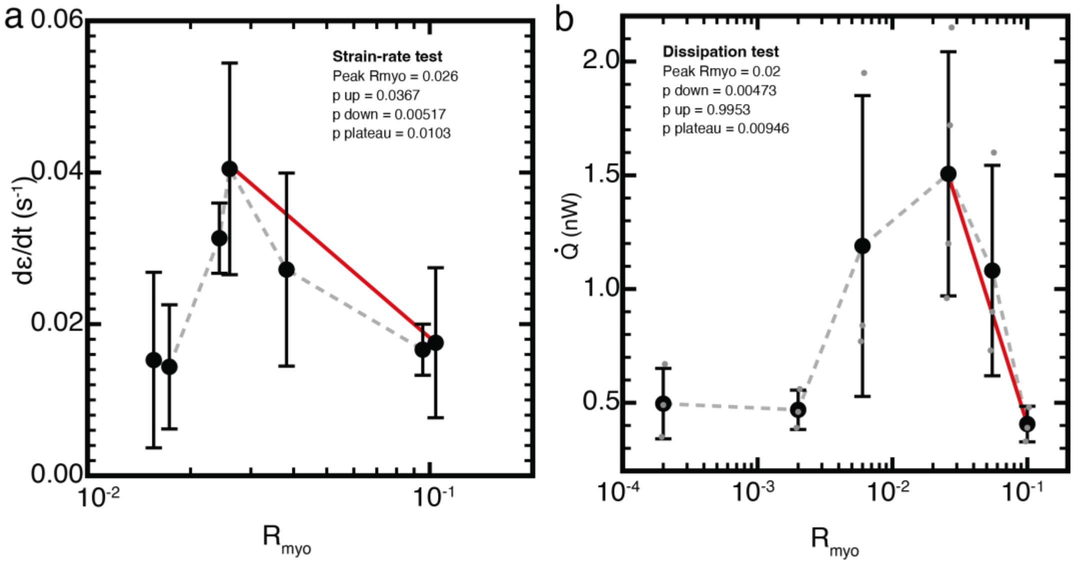

**SFig 1. Statistical test for monotonicity. a)** Strain rate, dε/dt, plotted as a function of myosin concentration ratio, $R_{myo}$. Data are shown as mean ± standard deviation. The gray dashed line connects the mean values. The red line shows the fitted post-peak trend after the maximum at $R_{myo}$= 0.026. The post-peak strain-rate trend significantly decreased with increasing $R_{myo}$ ($p_{down}$ = 0.00517) and rejected a plateau/no-slope relationship ($p_{plateau}$= 0.0103). **b)** Heat dissipation rate, Q̇, plotted as a function of R_myo. Gray points show individual measurements, black points show mean ± standard deviation, and the gray dashed line connects the mean values. The red line shows the fitted (linear model least squares regression) post-peak trend after the maximum at $R_{myo}$= 0.02. Post-peak dissipation significantly decreased with increasing $R_{myo}$ ($p_{down}$ = 0.00473) and rejected a plateau/no-slope relationship ($p_{plateau}$ = 0.00946).

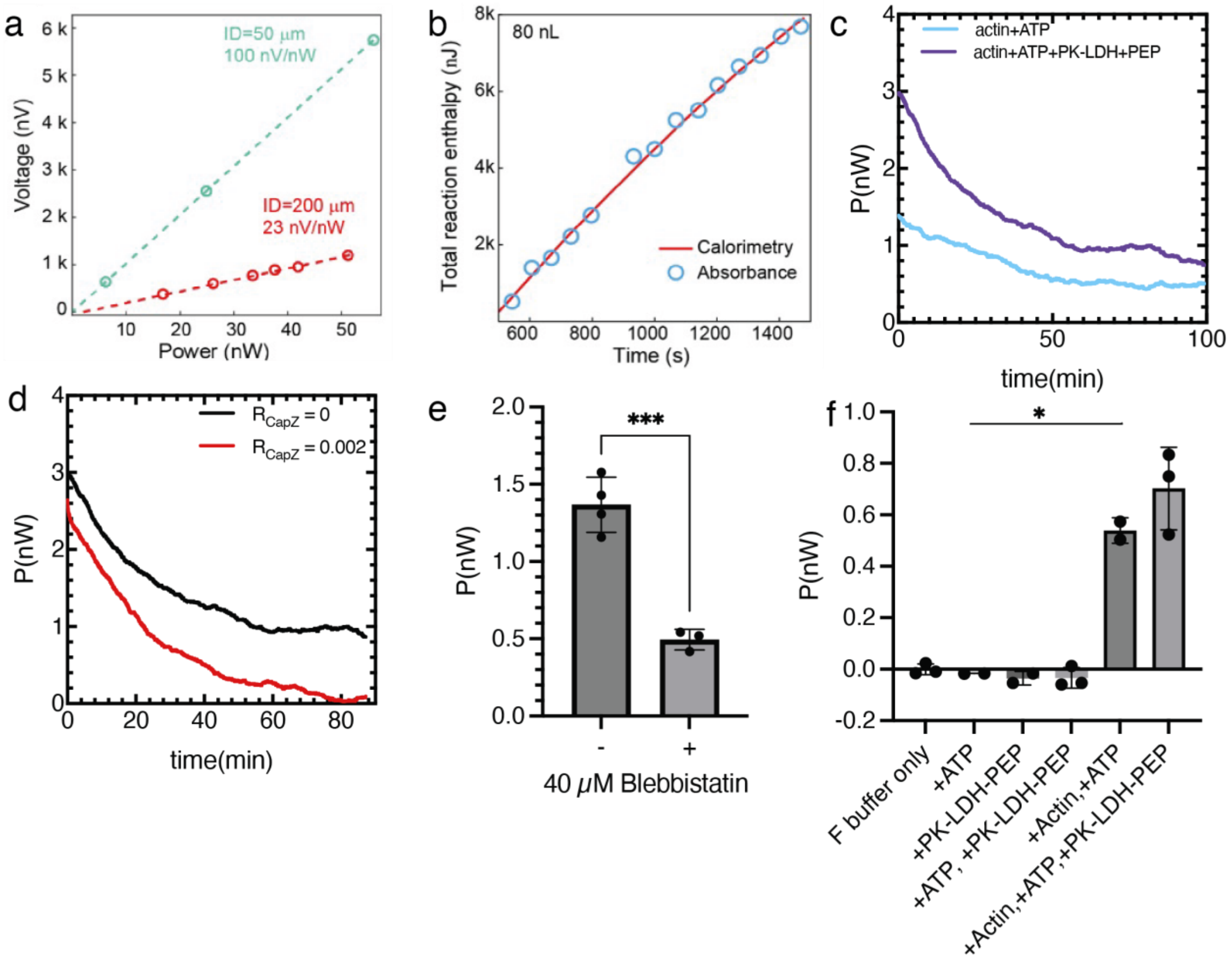

**SFig 2. Actomyosin heat dissipation control experiments.** a) Electrical calibration: Thermopile voltage response as a function of power applied to the on-chip heating element for sensors assembled with 50 μm-ID and 200 μm-ID capillaries. The 200 μm-ID configuration was used in this work of Zheng et al., *PNAS*, 2026[4]. b) Chemical calibration: Total reaction enthalpy measured calorimetrically compared with values obtained from NADH absorbance measurements for an 80 nL reaction volume[4]. c) Heat dissipation rate P as a function of time for actin polymerization with ATP only (blue) and actin with ATP + ATP regeneration (purple). d) Heat dissipation rate P as a function of time for actin polymerization with (red) and without (black) capping protein CapZ. e) Steady state heat dissipation rate P of actomyosin system without (-) and with (+) blebbistatin. N = 4 and 3 respectively. $p = 0.0008$. f) Steady state heat dissipation rate P of various control cases. N = 3,2,2,3,2,3 respectively. . $p = 0.0390$.

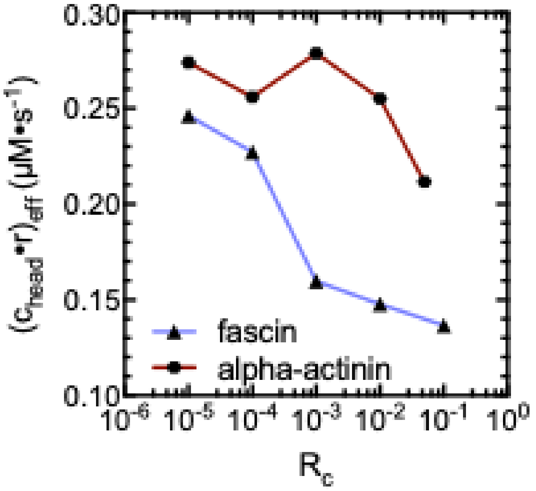

**SFig 3. Effective unbinding rate plot.** The effective unbinding rate for both crosslinker types under various $R_{CL}$ conditions. All conditions have $R_{myo}$ = 0.02. (Therefore $c_{head}$ should approximately be the same)

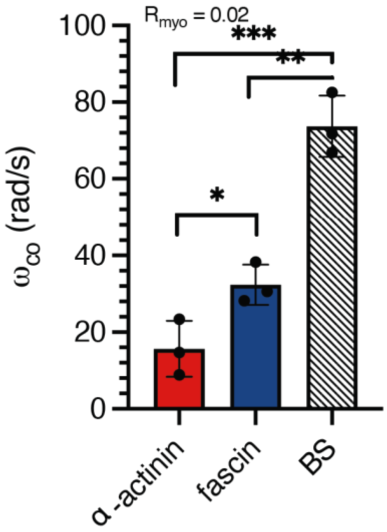

**SFig 4. Crossover frequency.** Crossover frequency $\omega_{co}$ for the three conditions in (c). N = 3 independent experiments for each condition. Welch's t-test is performed, $p_{f\text{-}a}$ = 0.0372, $p_{BS\text{-}a}$ = 0.0008, $p_{BS\text{-}f}$ = 0.003.

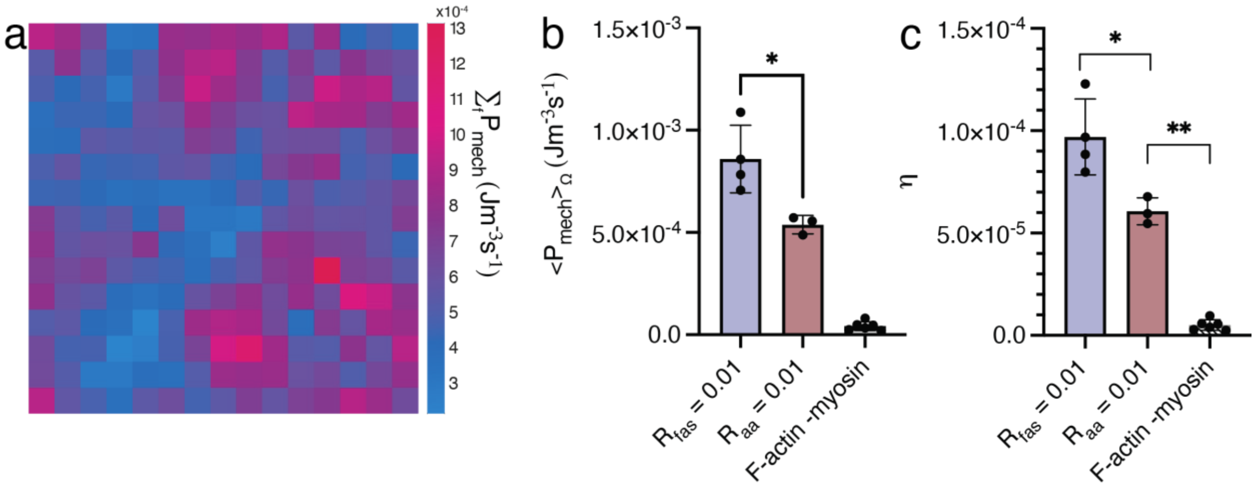

**SFig 5. Mechanical dissipation and efficiency plot.** a) Spatial map of the mechanical dissipation of the network summed over all frequencies for every window size. b) Averaged mechanical dissipation for $R_{fas}$ and $R_{aa}$ = 0.01, $R_{myo}$ = 0.02, and F-actin without myosin case. c) Mechanical efficiency $\eta$ for the three conditions. For b) and c), each point is an independent experimental run, the bars indicate the mean values and error bars are the s.t.d. of the mean. N = 4,3,6 respectively. For b), $p_{fas-aa}$ = 0.0251, and for c), $p_{fas-aa}$ = 0.0232, $p_{aa-actin}$ = 0.0027.

#### Supplementary movie captions

**Supplementary Movie 1. 2D Crosslinked actin network contraction under myosin active stress.** Fluorescent actin channel of fascin- (left) and α-actinin- (right) crosslinked network ($R_{fas}$ and $R_{\alpha a} = 0.2$) contracting into astors and rupturing under myosin active stress. Scale bar is 50 µm.

**Supplementary Movie 2. 3D fascin-crosslinked actomyosin network local deformation.** Fascin-crosslinked ($R_{fas}$ = 0.1) actin network exhibits local deformation overtime under myosin active stress. Scale bar is 10 µm.
